# Comparative metabolomics and microbiome analysis of Ethanol vs. OMNImet/gene®•GUT fecal stabilization

**DOI:** 10.1101/2023.08.29.555241

**Authors:** Heidi Isokääntä, Lucas Pinto da Silva, Naama Karu, Teemu Kallonen, Anna-Katariina Aatsinki, Thomas Hankemeier, Leyla Schimmel, Edgar Diaz, Tuulia Hyötyläinen, Pieter C. Dorrestein, Rob Knight, Matej Orešič, Rima Kaddurah-Daouk, Alex M. Dickens, Santosh Lamichhane, Alzheimer Gut Microbiome Project

## Abstract

Metabolites from feces provide important insights into the functionality of the gut microbiome. As immediate freezing is not always feasible in gut microbiome studies, there is a need for sampling protocols that provide stability of the fecal metabolome and microbiome at room temperature (RT). For this purpose, we investigated the stability of various metabolites and the microbiome (16S ribosomal RNA) in feces collected in 95% ethanol (EtOH) or OMNImet®•GUT/ OMNIgene®•GUT. To simulate in field-collection scenarios, the samples were stored at different temperatures at varying durations (24h +4°C, 24h RT, 36h RT, 48h RT, and 7 days RT), and compared to aliquots immediately frozen at -80°C. We applied several targeted and untargeted metabolomics platforms to measure lipids, polar untargeted metabolites, endocannabinoids, short chain fatty acids (SCFAs), and bile acids (BAs). We found that SCFAs in the non-stabilized samples increased over time, while a stable profile was recorded in sample aliquots stored in 95% EtOH and OMNImet®•GUT. When comparing the metabolite levels between fecal aliquots stored at room temperature and at +4°C, we detected several changes in microbial metabolites, including multiple BAs and SCFAs. Taken together, we found that storing fecal samples at room temperature and stabilizing them in 95% EtOH yielded metabolomic results comparable to flash freezing. We also found that overall composition of the gut microbiome did not vary significantly between different storage types. However, there were notable differences observed in alpha diversity. Taken together, the stability of the metabolome and microbiome in 95 % EtOH provided similar results as the validated commercial collection kits OMNImet®•GUT and OMNIgene®•GUT, respectively.

**IMPORTANCE:** The analysis of the gut metabolome and microbiome requires the separate collection of fecal specimens using conventional methods or commercial kits. However, these approaches can potentially introduce sampling errors and biases. In addition, the logistical requirements of studying large human cohorts have driven the need for home collection and transport of human fecal specimens at room temperature. By adopting a unified sampling approach at room temperature, we can enhance sampling convenience and practicality, leading to a more precise and comprehensive understanding of gut microbial function. However, the development and applications of such unified sampling systems still face limitations. The results presented in this study aim to address this knowledge gap by investigating the stability of metabolites and the microbiome (16S ribosomal RNA) from fecal samples collected using 95% EtOH, in comparison to well-established commercial collection kits for fecal metabolome (OMNImet®•GUT) and microbiome (OMNIgene® •GUT) profiling. Additionally, we perform a comparative analysis of various platforms and metabolomic coverage using matrices containing ethanol, evaluating aspects of sensitivity, robustness, and throughput.

## INTRODUCTION

The gut microbiome is considered an “essential organ” that contributes to the regulation of host development and physiology and facilitates host metabolism (1, 2) and is often linked to various human health conditions (3, 4), including inflammatory bowel disease (5), obesity (6), and multiple neurological disorders (7, 8). The interaction and dynamics between the host and gut microbiome are mediated by metabolites, which serve as vital signaling molecules (9). Integrated microbiome and metabolome analyses have emerged as the foremost promising approach to unveil host-microbiota interactions in the context of disease risk (10).

Over the last decade, fecal metabolomics has received increasing attention as fecal metabolites offer important insights into the functional aspects of the gut microbiome. The molecules associated with the gut microbiome also have the potential to be used in therapeutic strategies and biomarker discovery (11). Despite the growing interest, among the limiting factors in this field are practical challenges involved in the collection of human fecal samples. Albeit logistically impractical, the immediate homogenization and freezing of fecal samples at -80 °C is considered the gold standard for metabolite preservation because it halts enzymatic activity, hydrolysis, oxidation, and other degradation processes (12). Recently, studies of large human cohorts increased the demand for home collection of human fecal specimens, with the aim to reduce cost as well as to improve practicality, donor privacy, and convenience. Although home collection is a convenient option, it includes multiple steps that can involve temperature fluctuations. Moreover, storing and shipping frozen fecal samples can be inconvenient for participants and prohibitively expensive for researchers. This highlights the need to have room temperature storage facilities accessible (13). To address this need specifically for metabolomics analysis, available sampling devices such as DNA stabilization tubes and fecal immunochemical test tubes were critically assessed (14, 15). These studies found that the numerous detergents, buffers, salts, and other additives in the examined collection tubes deemed them inferior or unsuitable for analysis by liquid chromatography (LC) and mass spectrometry (MS). In addition, some of these collection kits can significantly distort the metabolic profile of feces samples compared to the gold standard of flash freezing (14, 15). A few studies tested 95% ethanol (EtOH) as a fecal sample preservative and found it suitable for metabolomics (10, 15). Ethanol prevents microbial growth and stabilizes the microbiome until profiling, while partly stabilizing the metabolome, as it prevents enzymatic metabolism by the fecal microbiota and affects chemical degradation processes. Another stability contributor effect of 95% EtOH is its frozen state at -80°C, hence no freeze-thaw cycles occur and disrupt the sample profile. Fecal collection and storage tubes containing 95% EtOH (OMNImet® •GUT, DNA Genotek, Canada) have been introduced as a kit tailored specifically for metabolomics analysis. A corresponding kit for the microbiome (OMNIgene® •GUT) has been used in several studies (16, 17). To our knowledge, only a few studies compared feces collection methods for simultaneous gut microbiota profiling and fecal metabolomics (10). Notably, studies comparing collection with 95% EtOH and commercial fecal collection kits are lacking.

Here we aim to fill the current knowledge gaps and examine the stability of metabolites and microbiome (16S ribosomal RNA) in feces samples collected in 95% EtOH compared to OMNImet® •GUT and OMNIgene® •GUT, the validated commercial collection kits for fecal metabolome and microbiome profiling. We also compare different platforms and metabolomic covers with EtOH -containing matrices in terms of sensitivity, robustness, and throughput.

## MATERIALS AND METHODS

### Studydesign

Stool samples were collected from four healthy human volunteers for metabolomics and microbiome profiling. For metabolomics, four different sample types were collected: immediately frozen feces, crude feces, feces in EtOH 95%, and commercially available OMNImet® •GUT kit that contains 95% EtOH. Each experimental condition had three replicates of each sample for metabolomics. The remaining samples were used for microbiome profiling. Before sampling, aliquot tubes were spiked with stability standards (Cholic acid-24-13C 2,5ppb, Palmitic acid (1-13C, 99%) 5ppm, Hippuric acid-d5 5ppm, Indole-2,4,5,6,7-d5-3-acetic acid 5ppm Nicotinamide-d4 (Major) 5ppm Sodium Butyrate-13C4 5ppm Sucrose-1-13Cfru 5ppm L-tyrosine-d4 5ppm Cortisol-1,2-d2 5ppm) and dried using a speedvac. Fecal samples were collected from 4 subjects in the morning next to the operating laboratory. Processing time with golden standard samples was 15-20 min from sampling to freezing. Three sub-tubes were obtained from different parts of the stool and homogenized with a spatula. These 3 homogenates represented replicates. Samples were divided into aliquots for different test designs (crude, 95% EtOH, OMNImet® •GUT, golden standard, RT/-80, 24h-7 days) with 3 replicates of each donor. Each aliquot had approximately 150 mg ±10mg stool in each prepared tube. Next, storage fluid was added at a ratio of 1:4 (600 ul of OMNImet® •GUT solvent or EtOH 95%) to tubes. For crude and golden standard samples, 100-ul ultra-pure water was added to make homogenization easier. All samples/aliquots were homogenized with a bullet homogenizer. Samples were kept at desired temperatures for the desired time and then stored in -80°C until analysis (1-5 months). For microbiome profiling, three sample types were collected: immediately frozen crude feces, feces in EtOH 95%, and commercially available OMNIgene®.GUT kit. Samples in 95% EtOH and OMNIgene®.GUT (including replicates mentioned above) were kept at RT for 24h, 48h, and 7 days and then stored in -80°C until DNA extraction (1-5 months).

### Analysis of fecal metabolome and microbiome

#### Metabolite extractions

We used a combined sample preparation method for SCFA, bile acids and untargeted metabolites. Prior to the extraction, samples were balanced by adding 100 ul water to EtOH/Omni-tubes and 600 µl EtOH to crudes and immediately frozen samples. Quality control sample (QC) was prepared by pooling an aliquot of 20 µl of each sample. The extraction included 100 µl of each sample and 1000 µl crash solvent (ethanol and Internal standards). After vortexing, samples were filtered through plates with vacuum. Filtrates were divided to 3 parts and stored at -80°C. Vials for BA analysis were dried under nitrogen, resuspended and stored at -80°C until analysis. Vials for untargeted metabolites were dried and stored at -80C before analysis. Filtrates for SCFA were stored on plates at -80°C. In this extraction we had following internal standards: Propionic acid-d6, Lithocholic acid (LCA)-d4, taurocholic acid (TCA)-d5, glycoursodeoxycholic acid (GUDCA) -d4, Glycocholic acid (GCA)-d4, Cholic acid (CA) -d4, Ursodeoxycholic acid (UDCA) -d4, glycochenodeoxycholic acid (GCDCA) -d4, Chenodeoxycholic acid (CDCA) -d4, Deoxycholic acid (DCA) -d4, Glycolithocholic acid (GLCA) -d4, Heptadecanoic acid, Deuterium-labeled Valine, Deterium-labeled Succinic acid, Deterium-labeled glutamic acid, MPFAC-MXA and d4-Androsterone.

For SCFAs, the sample aliquot was derivatized by adding 50 µL of 50 mM 3-NPH in 3:7 H2O: MeOH, followed by addition of 50 µL of 50 mM EDC in 3:7 H2O: MeOH, 50 µL Pyridine (7% in 3:7 H2O: MeOH). The mixture was then incubated for 60 min at room temperature, after which 100 µL Formic acid (0,2 % in 3:7 H2O: MeOH) was added to the mixture to quench the reaction. Second extraction was for lipidomics using liquid-liquid extraction. 10 µL of 0.9% NaCl was added to 10 µL of fecal sample (1:4) in EtOH. Internal standards in CHCl_3_: MeOH (2:1) were added 120 µl, (2.5 ppm). After vortexing, samples were let to stand on ice for 30 minutes. Then the samples were centrifuged (9400 × g, 5 min, 4°C). Next, 60 µL of the lower layer was collected and transferred to a LC vial with an insert and 60 µL of CHCl3: EtOH (2:1, v/v) was added. The samples were stored at -80°C until analysis. This extraction included following internal standards: PE(17:0/17:0), SM(d18:1/17:0), Cer(d18:1/17:0), PC (17:0/17:0), LPC (17:0), PC (16:0/d31/18:1), TG (17:0/17:0/17:0).

Third extraction was for endocannabinoids and it included 200 ul of fecal slurry and 400ul of crash solvent. To avoid contact with plastic, glass vials and syringes were utilized during extraction. Samples were vortexed and incubated for 30min in -20°C. And then filtered through plate. Filtrates were evaporated under a gentle stream of nitrogen +35°C and reconstituted with the final solvent 50ul/sample (40 % water, 30 % ACN and 30 % IPA) before run. This extraction included the following internal standards: THC-COOH-d9, 2-AG-d5, NADA-d8, AEA-d8 and AA-d8.

#### Metabolite analysis

SCFA and BAs were analyzed with an Exion LC system coupled with QTrap 5500 MS interfaced with a Turbo V electrospray ion source (SCIEX, Framingham, MA, USA). The lipids were analyzed with Exion LC system coupled with TripleTOF 6600 MS interfaced with a DuoSpray electrospray ion source (SCIEX, Framingham, MA, USA). The ECs were analyzed with an Exion LC system coupled with QTrap 7500 MS interfaced with an OptiFlow Pro electrospray ion source (SCIEX, Framingham, MA, USA). Untargeted metabolites were measured with gas chromatography and mass spectrometry (GS-TOF-MS).

For BA analysis, the chromatographic separation was carried out using an Acquity Premiere HSS T3 column (100 mm × 2.1 mm i.d., 1.8 µm particle size), fitted with a C18 precolumn (Waters Corporation, Wexford, Ireland). Mobile phase A consisted of water:methanol (v/v 70:30) and mobile phase B of methanol with both phases containing 2mM ammonium acetate as an ionization agent. The flow rate was set at 0.4 mL min-1 with the elution gradient as follows: 0-1.5 min, mobile phase B was increased from 5% to 30%; 1.5-4.5 min, mobile phase B increased to 70%; 4.5-7.5 min, mobile phase B increased to 100% and held for 5.5 min. A post-time of 5 min was used to regain the initial conditions for the next analysis. The total run time per sample was 18 min. The injection volume used was 5 µL. The analyses were performed in negative ion mode and Analyst v1.7.3 (AB SCIEX) was used for all data acquisition.

The SCFA was carried out on an Acquity UPLC BEH C18 column (2.1 x 100 mm, 1.7 µm; Waters, Milford, USA) using as mobile phase (A) 0.1 % formic acid in water and (B) 0.1 % formic acid in acetonitrile. Samples were eluted at 0.5 mL min-1, starting with 10% B and increasing to 100 % B in 10 min, then holding at 100% B for 2.1 min, returning to 10% B and holding for 2 min. Column temperature was maintained at 50 °C, while the auto sampler was maintained at 10 °C during analysis. The injection volume was 5 µL. The analyses were performed in negative ion mode and Analyst v1.7.3 (AB SCIEX) was used for all data acquisition.

The lipidomics analysis was carried out on an ACQUITY UPLC BEH C18 column (2.1 mm × 100 mm, particle size 1.7 μm) by Waters (Milford, USA) The eluent system consisted of (A) 10 mM ammonium acetate in H2O and 0.1% formic acid and (B) 10 mM ammonium acetate in ACN: IPA (1:1) and 0.1% formic acid. The gradient was as follows: 0-2 min, 35% solvent B; 2-7 min, 80% solvent B; 7-14 min 100% solvent B. The injection volume was 1 µL. All analyses were performed in positive ion mode and Analyst v1.8.1 (AB SCIEX) was used for all data acquisition.

The ECCs analysis was carried out on an XBridge BEH C18 column (2.1 x 150 mm, 2.5 µm; Waters, Milford, USA) using as mobile phase (A) 1mM ammonium acetate and 0.1 % formic acid in water and (B) 1mM ammonium acetate and 0.1 % formic acid in ACN: IPA (1:1). Samples were eluted at 0.4 mL min-1, starting with 60% B a holding for 0.3 min, then increasing to 100 % for 5 min and holding at 100% B for 2.5 min, returning to 10% in 0.1 min and holding for 3 min. Column temperature was maintained at 40 °C, while the autosampler was maintained at 15 °C during analysis. The injection volume was 10 µL. The analyses were performed in positive and negative ion mode and SCIEX OS v3.0 (AB SCIEX) was used for all data acquisition.

#### Gut microbiome analysis

Microbial DNA was extracted using a DNA Stool 200 Kit special H96 (PerkinElmer, Turku, Finland) kit with a corresponding Chemagic with Magnetic Separation Module I (MSM I) extraction robot. Microbial composition was determined by sequencing the V3V4 region of the 16S ribosomal gene using a MiSeq platform (Illumina, USA). The sequence library was constructed according to the Illumina library preparation protocol with minor differences to the V3V4 protocol. After PCR, 8µL of the integrity of the product was analyzed with 1,5% TAE agarose gel (120V, 1h). The concentration of the library samples was measured with a Qubit Fluorometer using a Qubit dsDNA High Sensitivity Assay kit. The 4 nM library pool was denatured, diluted to a concentration of 4pM and an 8% denaturalized PhiX control (Illumina, USA) was added. The library samples were sequenced with a MiSeq Reagent kit v3, 600 cycles (Illumina, USA) on a Miseq system with 2x 300 base pair (bp) paired ends following the manufacturer’s instructions. A positive control, Zymobiomics Microbial community DNA standard, (Zymo Research, USA)) and a negative control (PCR grade water) were included in library preparation to control the PCR. Lysis buffer was used as negative control in DNA extraction to control contamination.

#### Statistical analysis

For metabolomics data all multivariate statistical analyses were based on log-transformed intensity data. The transformed data were auto scaled prior to multivariate analysis to improve the global interpretability. To account for the inconsistency in the fecal water content the measured metabolites were normalized to the dry weight in the stool. The multivariate analysis was done using the PLS Toolbox 8.2.1 (Eigenvector Research Inc., Manson, WA, USA) in MATLAB 2017b (Mathworks, Inc., Natick, MA, USA). ANOVA-simultaneous component analysis (ASCA) a multivariate extension of ANOVA analysis was performed to allow interpretation of the variation induced by the different factors including time, individual, and collection matrix. Subsequently, for univariate analysis, the level of each metabolite in each sample storage matrix (i.e. crude feces without any solvent, feces in 95% EtOH, and feces in OMNImet® •GUT) was divided by the level of the same metabolite species in the paired immediately frozen sample (golden standard sample). For instance, the concentration of butyrate in the 95% EtOH was divided by the concentration of butyrate in the immediately frozen sample (golden standard). The fold difference was calculated by dividing the mean concentration of a given metabolite species in one group by another. The difference in the metabolites between the groups were tested using a multivariate linear model with using MaAsLin2 package in R, taking into account random effects within an individual sample or subject. The resulting nominal p-values were corrected for multiple comparisons using Benjamin and Hochberg approach. The adjusted p-values < 0.25 (q-values) were considered significantly different among the group of hypotheses tested.

Microbiome data analyses were performed in R Bioconductor ecosystem (R version 4.2.3) and with a CLC Microbial Genomics Module (CLC Genomics Workbench 21.0.3, Qiagen, USA), which complies with QIIME2 (18). Differences in beta diversity between methodological settings were evaluated with distance based redundancy analysis and PERMANOVA. We used Bray-Curtis dissimilarity. Moreover, beta diversity by Jaccard, UniFrac and Unweighted UniFrac were plotted. We calculated Shannon Index, number of observed OTUs, Chao1 and Simpson Index. All of the indices were used in the ICC testing. Differences in Shannon Index were assessed with two sample t-test assuming equal variance (Levene’s test).

## RESULTS

### Fecal metabolome and lipidome profiles

Targeted lipid profiling and untargeted metabolomics analysis were conducted on 168 fecal samples (aliquoted from four individuals) simulating different conditions of sample storage matrix (i) crude feces without any solvent, (ii) feces in 95% EtOH, and (iii) feces in OMNImet® •GUT. For each sampling condition, we obtained three biological replicates from different areas of a fecal sample (Fig. 1). To investigate the effects of storage time and temperature, we immediately froze corresponding aliquots of the homogenized fecal sample [i.e., 3 replicates X 4 donors] at -80°C (see Fig. 1), and then stored them for varying durations at different temperatures: 24 hours at +4°C (except OMNImet® •GUT), 24 hours at room temperature (RT), 36 hours at RT, 48 hours at RT, 48 hours at +4°C (except OMNImet® •GUT), and 7 days at RT.

**Figure 1.**
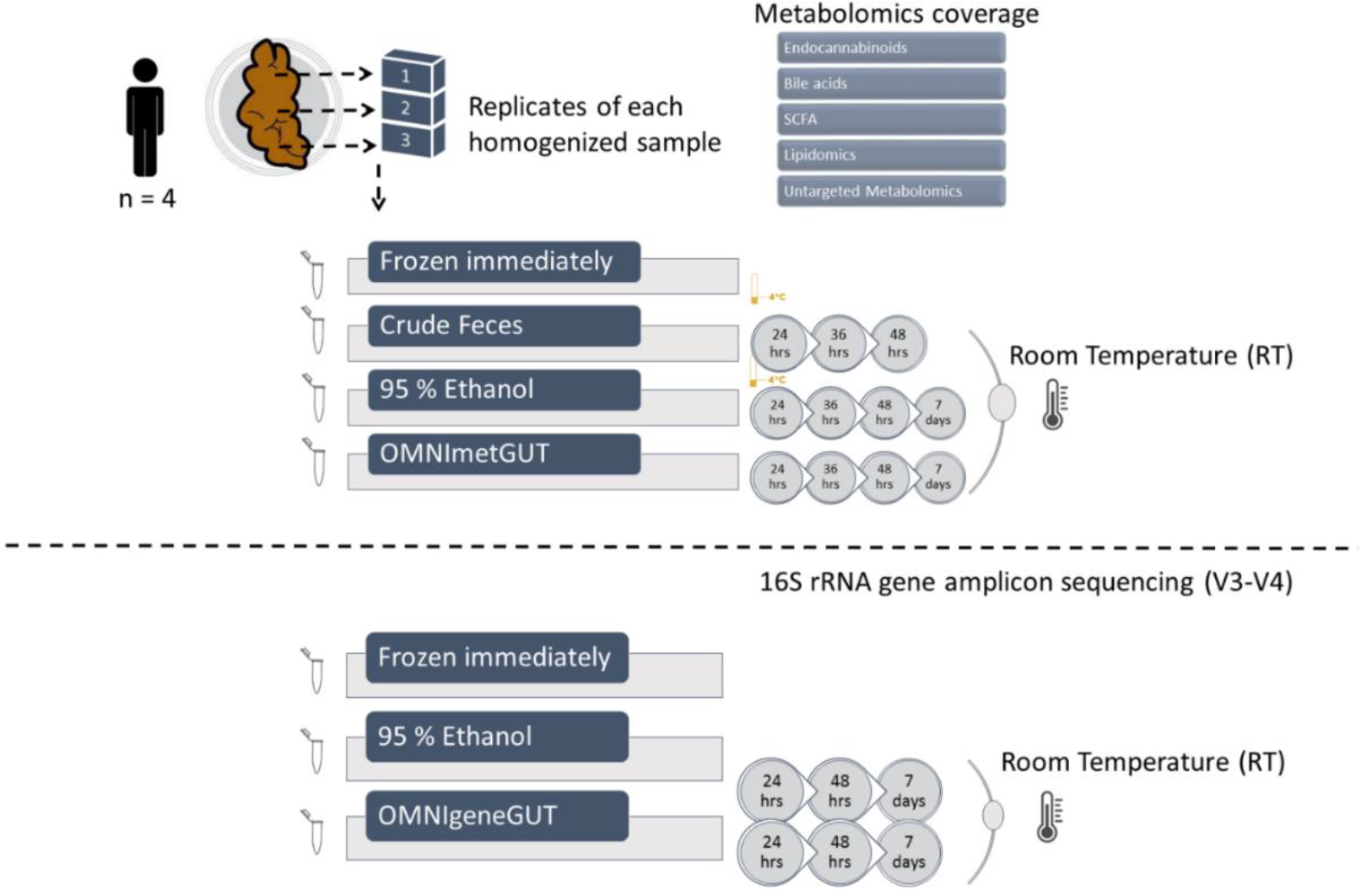
An overview of the study design is presented, illustrating the feces samples collected for metabolite measurement, as well as the number of matched feces samples for 16S rRNA gene sequencing at each time point. In this study, we analyzed aliquots of feces samples obtained from the fecal sample that were immediately frozen at -80°C and stored for 24 h, 48 h, and 7 days at room temperature in 95% EtOH and OMNIgene®•GUT/OMNImet®•GUT. Additionally, the figure shows the coverage of targeted and untargeted metabolomics performed in this study.

Targeted analysis of lipids provided coverage of the following lipid classes: acylcarnitine (AC), cholesterol esters (CE), ceramides (Cer), diacylglycerols (DG), lysophosphatidylcholines (LPC), phosphatidylcholines (PC), sphingomyelins (SM) and triacylglycerols (TG). The metabolomics analysis included targeted short chain fatty acid (SCFA, n=7) and bile acids (BAs, n=33) including both primary (glycine/taurine conjugates) and secondary BAs as well as endocannabinoids (ECCs, n=9). These ECCs included palmitoylethanolamide (PEA), arachidonoyl glycerol (AG), 2-arachidonoyl glycerol ether (2-AGE), arachidonoyl ethanolamide (AEA), oleoylethanolamide (OEA), stearoyl ethanolamide (SEA), docosahexaenoyl ethanolamide (DEA), alpha-linolenoyl ethanolamide (aLEA), and arachidonic acid (AA). Untargeted metabolomics analysis provided coverage of the following classes: amino acids (regular amino acids and branched chain amino acids), carboxylic acids (mainly free fatty acids and other organic acids), hydroxy acids, phenolic compounds, alcohols and sugar derivatives.

#### Fecal metabolic profile

##### I. Multi-factorial analysis

To understand the contributions of different sampling factors to the fecal metabolome, we performed analysis of variance (ANOVA)-simultaneous component analysis (ASCA) with the following factors: subjects; sample storage matrix (crude, 95% EtOH, OMNImet® •GUT, immediately frozen); time (24 h, 36 h, 48 h, 7 days); temperature (RT, +4°C). We found that inter-individual differences (71.15, 65.1, 77.7, p = 0.0010) and subsequent sample storage matrix (8.2, 9.6, 2.5, p = 0.0010) had the strongest effect on BAs, SCFAs, and ECCs fecal profiles. In comparison, the ASCA could not find significant effects of the duration of storage (2.1, 2.0, 1.7, p > 0.05) and storage temperature (1.60, 1.09, 1.34, p > 0.05). Figure 2 illustrates the clustering differences in subject (Fig. 2a) and sample storage matrix (Fig. 2b) explained by the levels of fecal BAs. Similar trends were observed for SCFAs and ECCs (Supplementary Fig. S1 and Fig. S2). Similar analysis utilizing untargeted lipidomics and polar metabolites (see Supplementary Fig. S3), found only a minor inter-subject significant effect (15.3, 19.9, p = 0.0010), with no contribution of the sample storage matrix, storage time and temperature (p > 0.05). This may be attributed to larger within-group variance.

**Figure 2.**
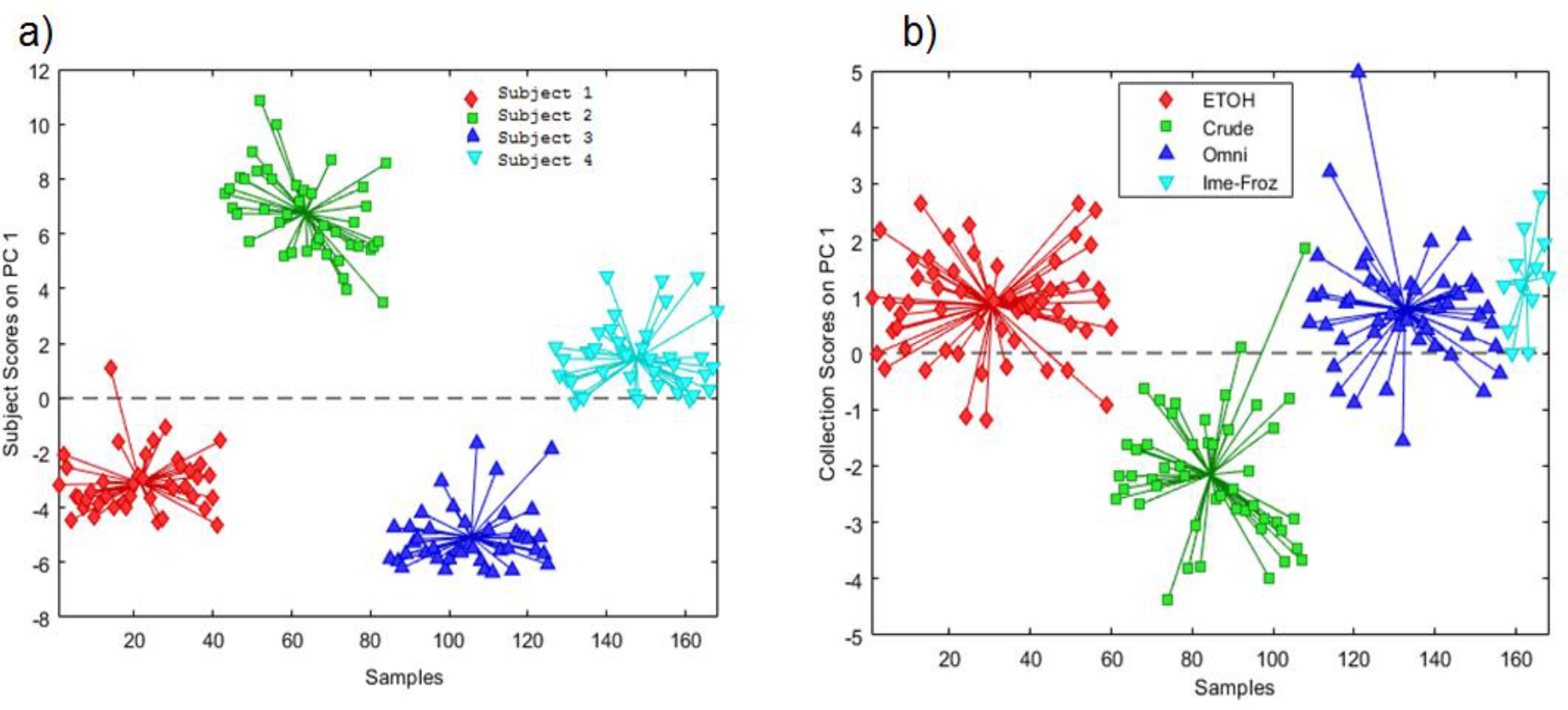
Principal component analysis (PCA) score plots based on ANOVA-simultaneous component analysis (ASCA). a) Principal component (PC1) score plot obtained based on inter-individual score in ASCA analysis. This figure represents the bile acid dataset arranged according to inter-individual samples in the PCA score plot. Here each sample is represented by a point and colored according to the individual (red diamond: Subject1, green square: Subject 2, blue triangle up: Subject 3, cyan triangle down: Subject4). b) Principal component (PC1) score plot obtained based on sample storage matrix score in ASCA analysis. This figure represents the bile acid profile arranged according to inter-individual samples in the PCA score plot. Here each sample is represented by a point and colored according to the sample storage matrix (red diamond: 95% EtOH, green square: crude, blue triangle up: OMNImet®•GUT, cyan triangle down: immediately frozen).

##### II. Metabolic changes throughout storage at room temperature

To better understand the effects of sample storage duration (24h, 36h, 48h, 7 days) at room temperature on the fecal metabolome, we analyzed the stability of metabolites in aliquots stored without solvent (crude), with 95% EtOH, and with OMNImet® •GUT tube. For each sample matrix, we calculated the relative change of each metabolite compared to the gold standard (immediately frozen sample).

We found that SCFAs in the crude samples increased over time, while a stable profile was recorded in sample aliquots stored in 95% EtOH or OMNImet® •GUT tube. In particular, butyric acid, isobutyric acid, and valeric acid increased by >1.5 fold in raw feces left at RT from 24 hours till day 48 hour (Fig. 3 a & b, p.adj < 0.25, Supplementary Fig. S4), suggesting continuous microbial activity over time. We also observed an increase in the levels of butyric acid, isobutyric acid, and valeric acid by 1.72, 1.73, and 1.47-fold (FC), respectively, within the first 24 hours compared to the sample that was frozen immediately. However, we observed a stable pattern of these SCFAs in fecal samples collected with 95% EtOH and/or OMNImet GUT solvent (Fig. 3 and Supplementary Fig. S4).

**Figure 3.**
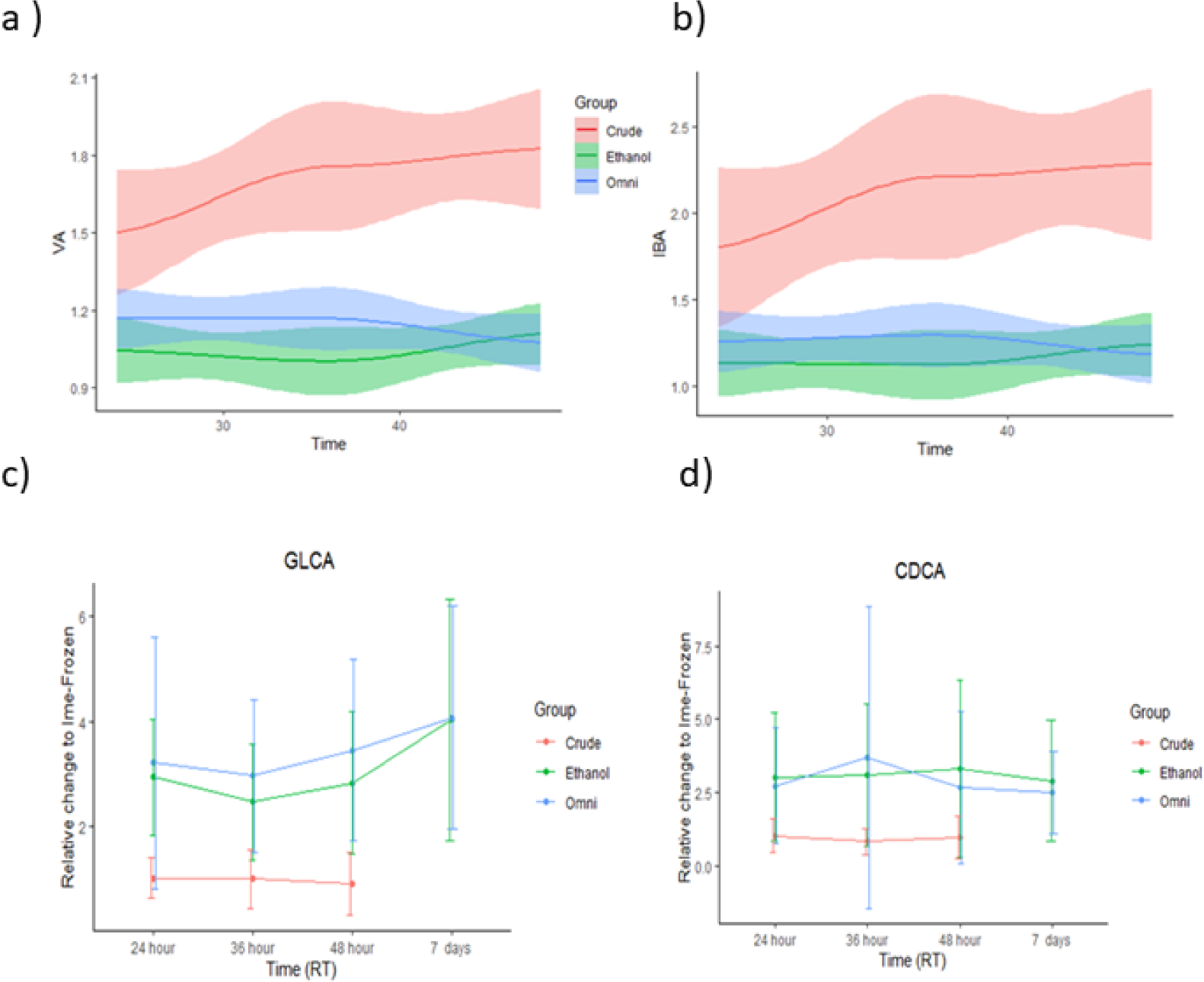
Alterations in metabolites during storage at room temperature. A loess curve plot showing the changes in the levels of two SCFA [(a) Valeric acid (b) Iso-butyric acid] over time (24h, 36h, 48h) in feces samples collected as crude, in 95% EtOH, and with OMNImet®•GUT solvent. X-axis shows the sample storage duration (24h, 36h, and 48h). Y-axis shows the relative change of each metabolite compared to the gold standard. VA is valeric acid and IBA is isobutyric acid. The changes in bile acids (BAs) over time (24h, 36h, 48h, 7 days) were examined in feces samples collected in three different ways: crude, in 95% EtOH, and in OMNImet®•GUT solvent. **c)** Glycolithocholic acid (GLCA) and d) Chenodeoxycholic acid (CDCA)

No time-dependent pattern was found along 7 days of storage at room temperature in all sample storage matrices (crude, 95% EtOH, OMNImet® •GUT, immediately frozen). We also analyzed the concentration differences of fecal bile acids, lipids, and untargeted metabolites over time in these sampling groups. Tauro- and/or glycoconjugated bile acids (GLCA, THDCA, GCDCA), lipids (mainly TGs), and unknown polar features increased over time (p < 0.05) and differed in at least one of the three sample storage matrices. However, none of these metabolites exceeded the significance level at the selected FDR threshold of 0.25 (Fig. 3 c & d, Supplementary Fig. S5-7, and Supplementary Table S1-S3).

##### III. Fecal Metabolic changes in 95% ethanol at different temperatures

We compared the differences of individual metabolite levels between fecal aliquots stored at room temperature and at +4°C. We found significant changes in microbial metabolites, including conjugated bile acids and SCFAs. GLCA content was 1.31 FC higher in samples stored at room temperature than in fecal aliquots stored at +4°C in 95% EtOH (Fig. 5). In crude feces, two BAs [7-oxo DCA (FC = 4.1), DHCA (FC = 0.45)] and five SCFAs, including acetic acid (FC = 1.51), butyric acid (FC = 1.64), isobutyric acid (FC = 1.37), valeric acid (FC = 1.28), propionic acid (FC = 1.32), and the internal stability standard Butyric acid-^13^C_4_(FC = 0.51) changed when stored at RT (Fig. 4, Supplementary Table 4-5). In contrast, these metabolites (e.g., SCFAs) remained stable in samples collected in 95% EtOH, which may be partly due to active enzymatic metabolism by the gut microbiota at room temperature but not when deactivated by 95% EtOH (Supplementary Table S4-S5).

**Figure 4.**
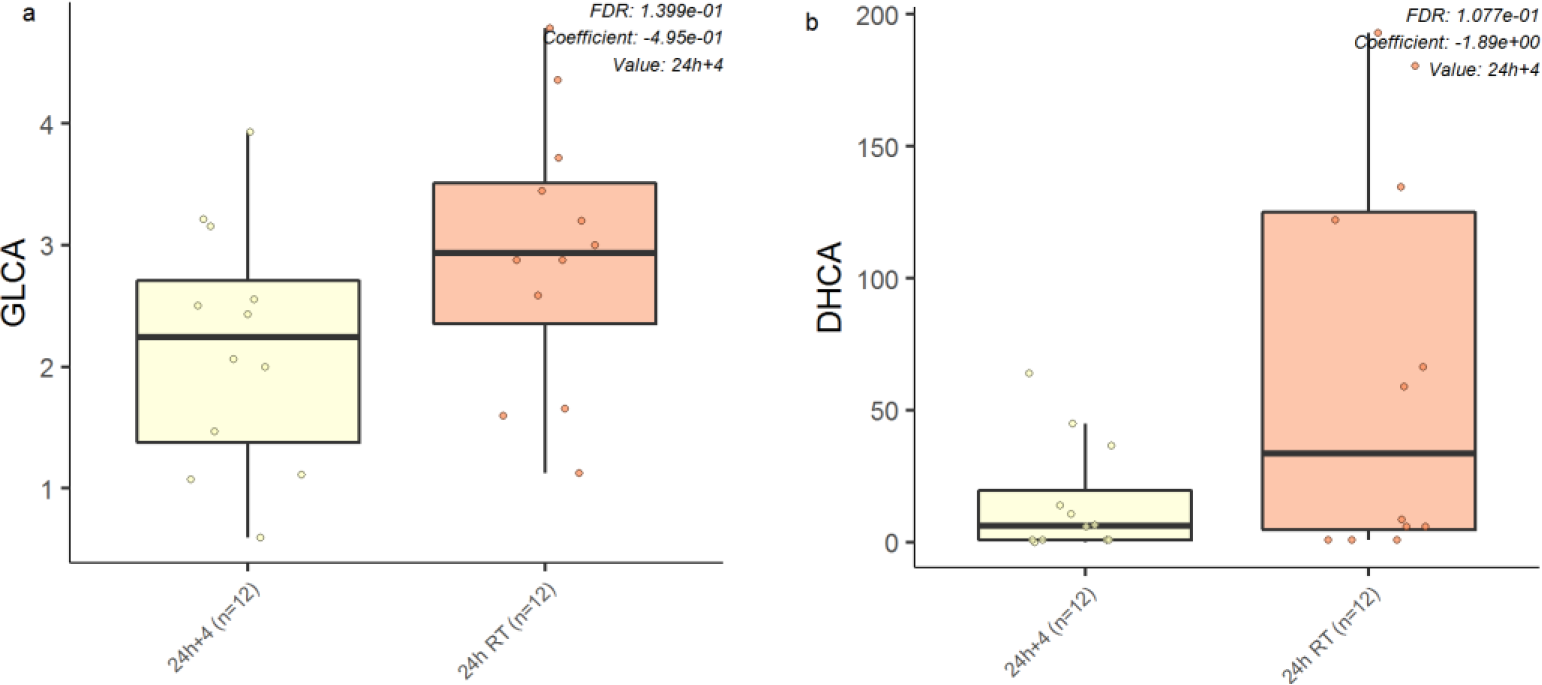
Stability of fecal metabolites at room temperature (RT) compared to +4°C for (a) 95% EtOH samples and (b) crude samples. Y axis denotes level of metabolites and X axis denotes the different temperatures

**Figure 5.**
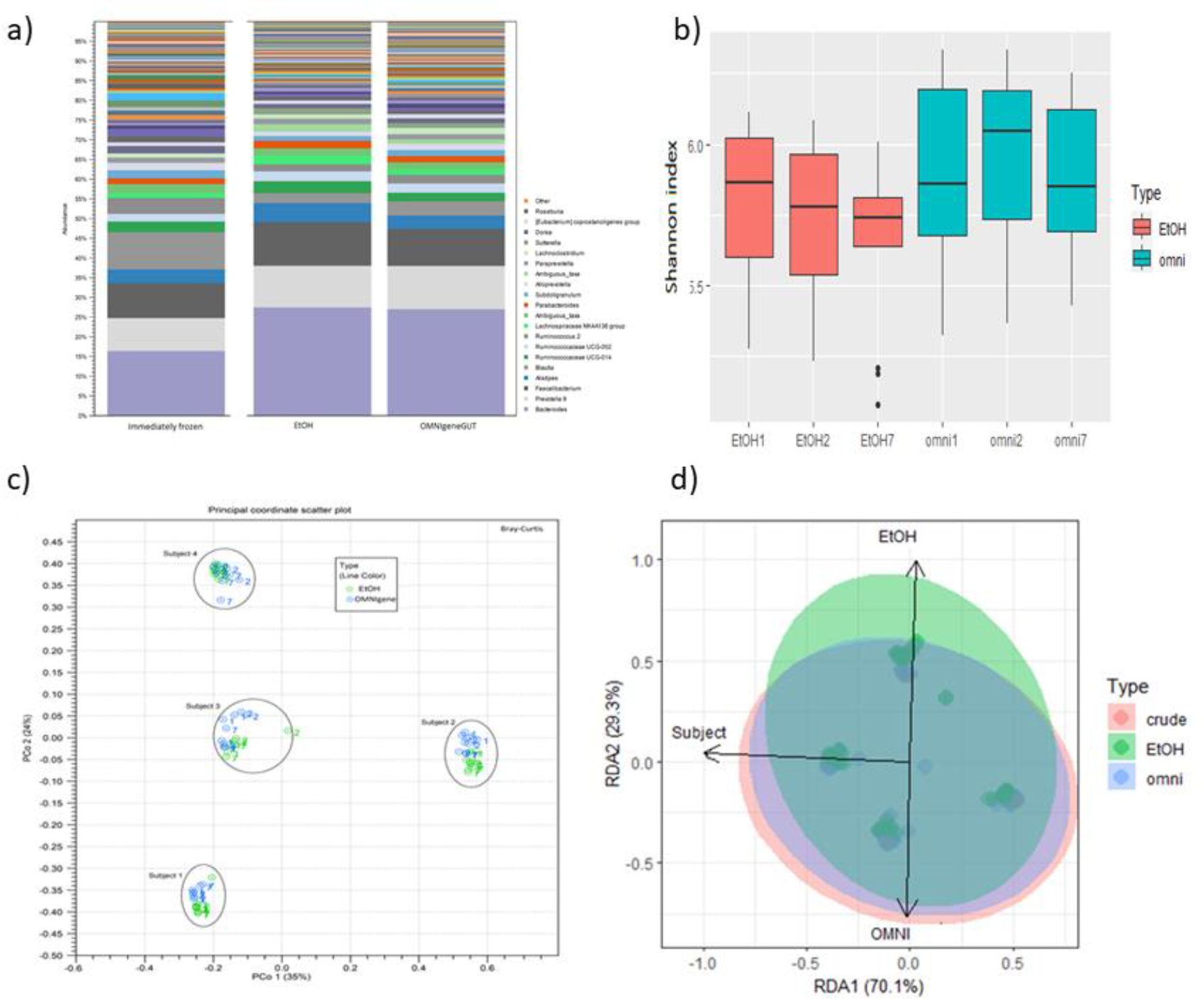
Stability of fecal microbiome in 95% EtOH stored at RT. a) Microbiome profiles by relative abundances in different storage types. Legend shows 20 most abundant genera. Main genera are the same, abundances differ between immediately frozen and samples in preservatives. b) Alpha diversity (Shannon) by storage type and time. Numbers indicate days of storage. c) Beta diversity by Bray-Curtis. A) Beta diversity by Principal coordinates analysis with Bray-Curtis dissimilarity metrics. Numbers indicate storage types with colors and storage days. Oval shapes form clusters of subjects. B) Distance-based redundancy analysis with Bray-Curtis representing dissimilarity between storage types with ellipses of 95% confidence interval.

#### Fecal microbiome profile

We aimed to determine the stability of the fecal microbiome in 95% EtOH stored at RT. The composition of the fecal microbiota was analyzed by 16S rRNA gene sequencing using V3-V4 hypervariable region (n = 118). These samples correspond to aliquots obtained from the fecal sample that was immediately frozen at -80°C and stored for 24 hours, 48 hours, and 7 days at RT in 95% EtOH and OMNIgene® •GUT (Fig. 1). The bacterial profiles with relative abundances from these collections are shown in Fig. 5a and are summarized for all subjects and time points. We found that the bacterial profiles of the fecal samples stored in 95% EtOH and OMNIgene® •GUT were similar. However, the immediately frozen samples differed from those stored in 95% EtOH and OMNIgene® •GUT mainly by Bacteroides and Blautia. Positive controls included in the NGS-protocol indicated high reproducibility, and the accuracy was sufficient based on the theoretical and identified abundances (Supplementary Fig. S8).

Similarly, we analyzed the differences in alpha diversity (Shannon index) among the three study groups. We found that inter-individual differences had a dominant effect on microbiome alpha diversity (Supplementary Fig. S9). Lower alpha diversity was observed in both storage solvents compared to immediately frozen samples. We also found significant decrease in alpha diversity in the longitudinal series of samples collected in 95% EtOH. Fig. 5b shows that alpha diversity was lower over time in 95% EtOH compared with OMNIgene (p=0.007). We also compared similarity between storage types using intraclass correlation coefficients with immediately frozen samples used as reference (ICC, see Supplementary Fig. S10). We compared similarity in alpha diversity metrics (Shannon, Simpson, Chao1, number of observed OTUs) and three most prevalent genera (*Bacteroides, Bifidobacterium, Faecalibacterium*) between storage types. OMNIgene®•GUT showed higher alpha diversity intra-class similarity with the immediately frozen samples than in the 95% EtOH samples.

Additionally, we also analyzed the differences in overall gut microbiota composition, i.e. beta diversity between samples collected in 95% EtOH and OMNIgene® •GUT. PcoA with Bray-Curtis showed little difference between storage types and time points (Fig. 5c). PcoA with Jaccard, UniFrac and Unweighted UniFrac showed parallel results, however there were less differences with UniFrac metrics (Supplementary Fig. S11-12). However, the score plot revealed that inter-individual variability is the key factor shaping the fecal microbiome (Fig. 5d). There were no significant differences in beta diversity between storage types (distance-based redundancy analysis with Bray-Curtis dissimilarity, PERMANOVA, p=0.3). However, inter-individual differences were significant (p= 0.001) as expected.

## DISCUSSION

The aim of the presented study was to establish the validity of an off-site fecal collection method that preserves the integrity of both the metabolome and the microbiome during freight and until processing. For this goal, 95% ethanol was selected as a preservative, and compared with OMNImet® •GUT and OMNIgene® •GUT, which are validated commercial collection kits for fecal metabolome and microbiome profiling, respectively. To our knowledge, this is the first study to directly compare the performance of the OMNImet® •GUT kit, which is designed to preserve fecal metabolites at room temperature, to flash freezing feces or using 95% ethanol. Unlike similar studies, we also utilized internal standards and biological replicates to assess the stability of detected metabolites at room temperature. We applied a wide range of metabolomics platforms: targeted metabolomics to measure SCFA, BAs, and endocannabinoids, as well as untargeted metabolomics to detect lipids and polar metabolites. Various studies have shown that both the human gut microbiota and metabolome are highly heterogeneous between individuals (19-21) . Indeed, our work showed that variance between fecal samples was mainly attributed to inter-individual differences and affected by the sample storage matrix, while less affected by the duration at room temperature when stored in in 95% EtOH and OMNImet® •GUT liquid.

We generally observed good metabolite stability in the 95% EtOH and OMNImet® •GUT kit, irrespective of the storage temperature. Our findings are consistent with previous studies suggesting that fecal samples in 95% EtOH or OMNImet® •GUT have comparable metabolite profiles to samples that were frozen shortly after collection (10, 13, 15, 22). In addition, 95% EtOH was reported to be the most suitable matrix for preserving the fecal metabolite profile in comparison to clinically used fecal collection kits that do not target metabolites (RNAlater, OMNIgene®•GUT, fecal occult blood test (FOBT) cards) (15, 22). Feces collected without any solvent was suitable for metabolomics analysis when frozen immediately after collection (15). However, it is worth noting that this approach is inconvenient, expensive, and not feasible for population-based studies on a large scale. Liu *et al*. demonstrated that metabolite measures obtained from OMNIgene® •GUT were comparable to those obtained from samples that were immediately frozen after collection for up to 21 days. These findings suggest that OMNIgene® •GUT is sufficient for obtaining data on the gut microbiome and gut metabolome (14). On the contrary, other studies report that OMNIgene® •GUT may not be optimal for collecting fecal samples for metabolomics profiles (10, 22).Clearly, more systematic studies are needed to examine the complementary single-sampling method, including OMNIgene® •GUT.

Research investigating the gut microbiome-metabolome relationship in health and disease has predominantly concentrated on water-soluble polar metabolites, with microbe-linked lipids receiving less emphasis (23-25). To aid in this effort, apart from the methodological comparison, our results also highlight that the fecal lipids can serve as a functional indicator of gut microbial metabolism.

Crude feces (stored without any solvent) at room temperature showed at least 50% increase in SCFA over 24-48 hours. SCFAs are primarily produced by gut microbes through the saccharolytic fermentation of complex microbiota-accessible carbohydrates (MACs) (26, 27)). Therefore, our results suggest that crude samples at room temperature are susceptible to microbial metabolism by specific microorganisms encoding MAC-degrading enzymes. This is corroborated by the increased SCFA levels we recorded at room temperature when compared to samples stored at 4°C. These results support the hypothesis that 95% EtOH can stabilize microbial activity, such as saccharolytic fermentation, even at room temperature that can prevent and reduce metabolite degradation. However, there were exceptions of conjugated BAs such as GLCA, GCDCA, GDCA, GCA, and several unknown metabolites (Supplementary Table S5) that appeared to be affected by temperature (nominal p-value < 0.05) and require further investigation.

We acknowledge that there are some limitations that need to be considered. The main limitation is that instead of collecting directly within each inspected matrix, we only simulated such collection, after processing a sample of collected crude feces. This is an inherent technical limitation of the study design that requires aliquots of the same biological sample to be compared across various collection matrices and storage conditions. Another potential limitation is the collection by trained team. For instance, using non-commercial 95% ethanol kits for sampling may present practical challenges compared to using commercial kits such as OMNIgene® •GUT. In current study, trained staff conducted the sampling; however, the effectiveness of using 95% EtoH for lay study participants for home collection needs to be tested. Notwithstanding this, ongoing Alzheimer’s Gut Microbiome Project (https://alzheimergut.org/) has already tested the effectiveness of using 95% EtoH. It is also worth noting that all subjects in this study were adults, and the metabolite and microbiome content of feces differ in different age ranges, such as in children. Therefore, further validation may be necessary to test fecal samples collected from a wider range of age groups. Another factor that limits the reliability of fecal sample collection is the variability in the volume and weight of the feces (water and fiber content). To address this, in the current setting we collected fecal samples volumetrically, using a weight-to-volume ratio of 1:4 (weight of feces to volume of ethanol 95%) to match the W:V ratio in the commercially available kit OMNIgene® •GUT. Notwithstanding that, two steps may be taken to address this issue. First, weigh the collection tube (95% ethanol kits) before sending it to the participant to obtain an accurate measurement of the fecal sample weight. Second, participants report their stool consistency, which is strongly associated with the composition of the gut microbiota and metabolite content. By considering dry weight we may be able to account for the physicochemical bias during the sample collection process. In terms of microbiome profiling, our study was limited to metataxonomics (*i*.*e*., 16S rRNA gene sequencing) analyses, and future studies should consider metagenomics (whole shotgun metagenomic sequencing) and study if 95% EtOH collection is suitable for metatranscriptomics (gene expression study).

## CONCLUSIONS

Overall, our study found that storing feces samples at room temperature and stabilizing them in OMNImet® •GUT or 95% EtOH yielded metabolomic results generally comparable with flash freezing. Specifically, we observed similar identities and abundances of detected biochemicals, as well as comparable metabolic profiles of the study subjects. Moreover, we characterized metabolic changes in crude feces over time, which could be attributed to microbiota activity and non-enzymatic reactions such as oxidation-reduction. Utilising 95% EtOH as fecal collection matrix can offer a more convenient and cost-effective way to collect and store feces samples at home. Individual differences in microbiome overall composition dominated those of storage type. However, OMNIgene® •GUT was slightly better than 95% EtOH at preserving microbiota based on alpha diversity. Further exploration of an existing commercial kit is ongoing and will expend the microbiome and metabolome assessment.

## Acknowledgement

We thank the Turku Metabolomics Center for the assistance and resources in the analyses of fecal metabolome and lipidome. This study was supported by the National Institute on Health grant (U19AG063744; PI: Kaddurdah-Daouk) and the Academy of Finland project grant, (No. 323171 to S.L.), (No. 333981 to M.O.), Swedish Research Council (grant no. 2016-05176 to T.H and M.O), Formas (grant no. 2019-00869 to T.H and M.O), and the Novo Nordisk Foundation (Grant no. NNF20OC0063971 to T.H. and M.O.). Further support was received by “Inflammation in human early life: targeting impacts on life-course health” (INITIALISE) consortium funded by the Horizon Europe Program of the European Union under Grant Agreement 101094099 (to M.O. and T.H.,) and Alzheimer’s Gut Microbiome Project (https://alzheimergut.org/).

Funding sources had no role in study design, the collection, analysis and interpretation of data, the writing of the report, or in the decision to submit the article for publication. All authors approved the final version and had final responsibility for the decision to submit for publication.

## Declarations

Dr. Kaddurah-Daouk is an inventor on a series of patents on the use of metabolomics for the diagnosis and treatment of CNS diseases and holds equity in Metabolon Inc., Chymia LLC and PsyProtix. All authors declare no competing interests.

## Ethics declarations

Fecal samples were obtained from anonymous fecal donations of four healthy individuals; informed consent was obtained from all donors. The Finnish legislation and ethical committee in University of Turku on Research involving Human Beings do not restrict research using anonymized biological material. An ethical review statement was not necessary because samples were given voluntarily with supporting information, no exceptionally strong stimulus was done, personal data was not used, fecal sampling is non-invasive, and the research does not involve a threat to the safety or harm or limit of a normal daily life to the participants or researchers.

