## Supplementaryfigures for "Comparative metabolomics and microbiome analysis of Ethanol vs. OMNImet/gene®•GUT fecal stabilization"

**Isokääntä *et al.***

### SUPPLEMENTARY MATERIALS

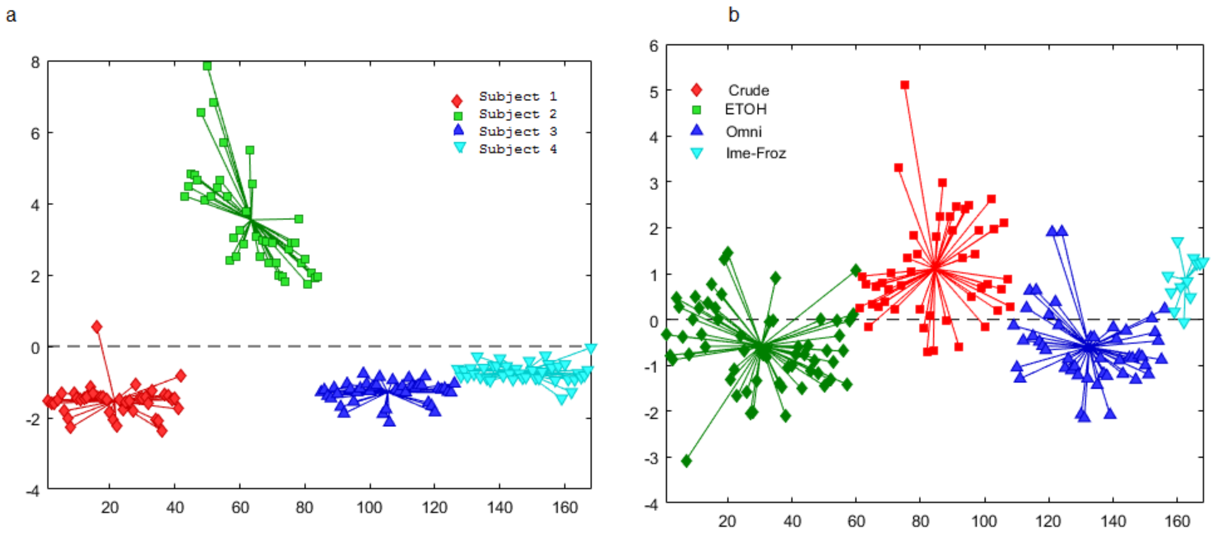

**Supplementary FigureS1.** For SCFA. Principal component analysis (PCA) score plots based on ANOVA-simultaneous component analysis (ASCA).

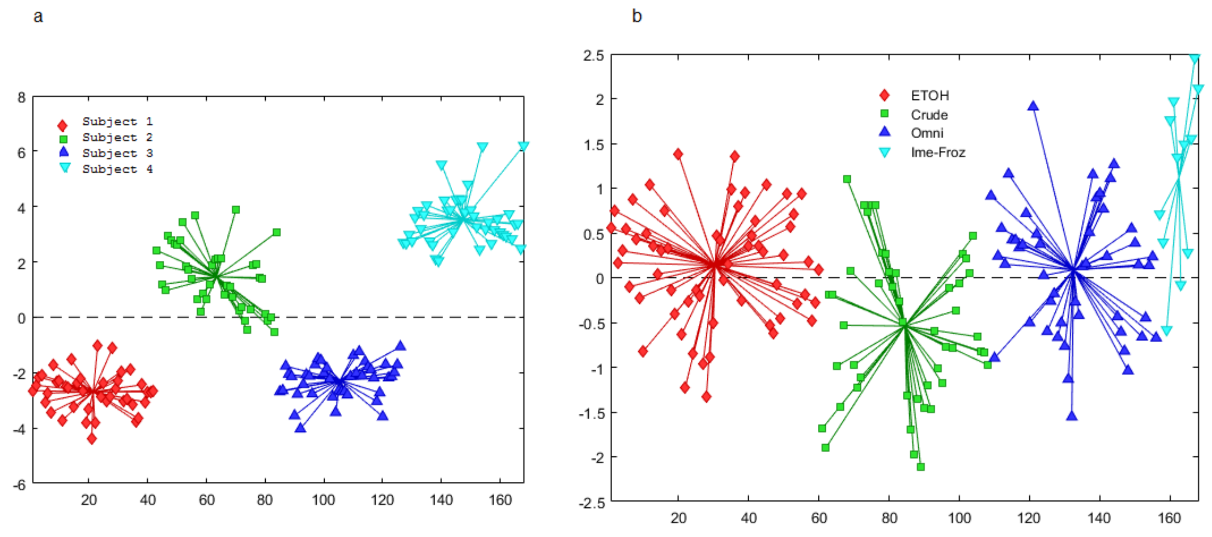

**Supplementary FigureS2.** For ECCs. Principal component analysis (PCA) score plots based on ANOVA-simultaneous component analysis (ASCA).

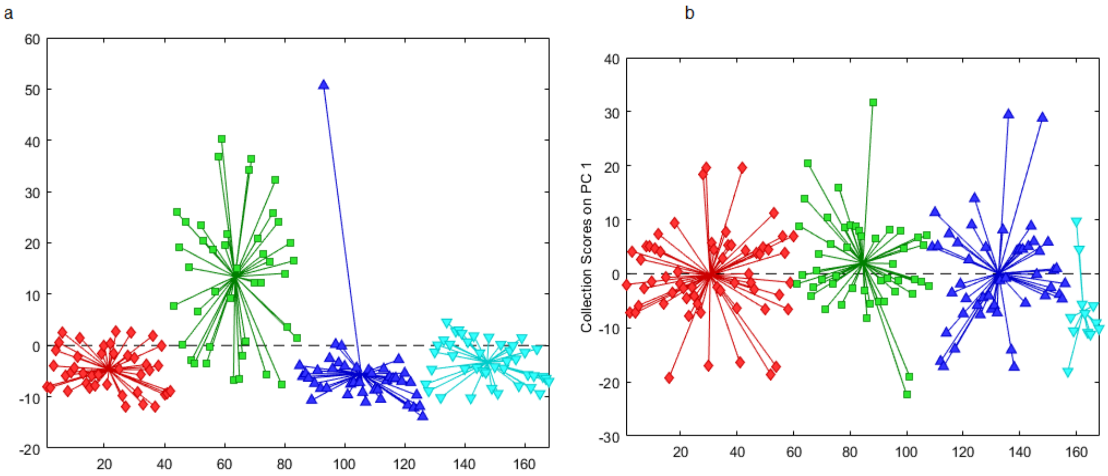

**Supplementary FigureS3.** For Lipids. Principal component analysis (PCA) score plots based on ANOVA-simultaneous component analysis (ASCA).

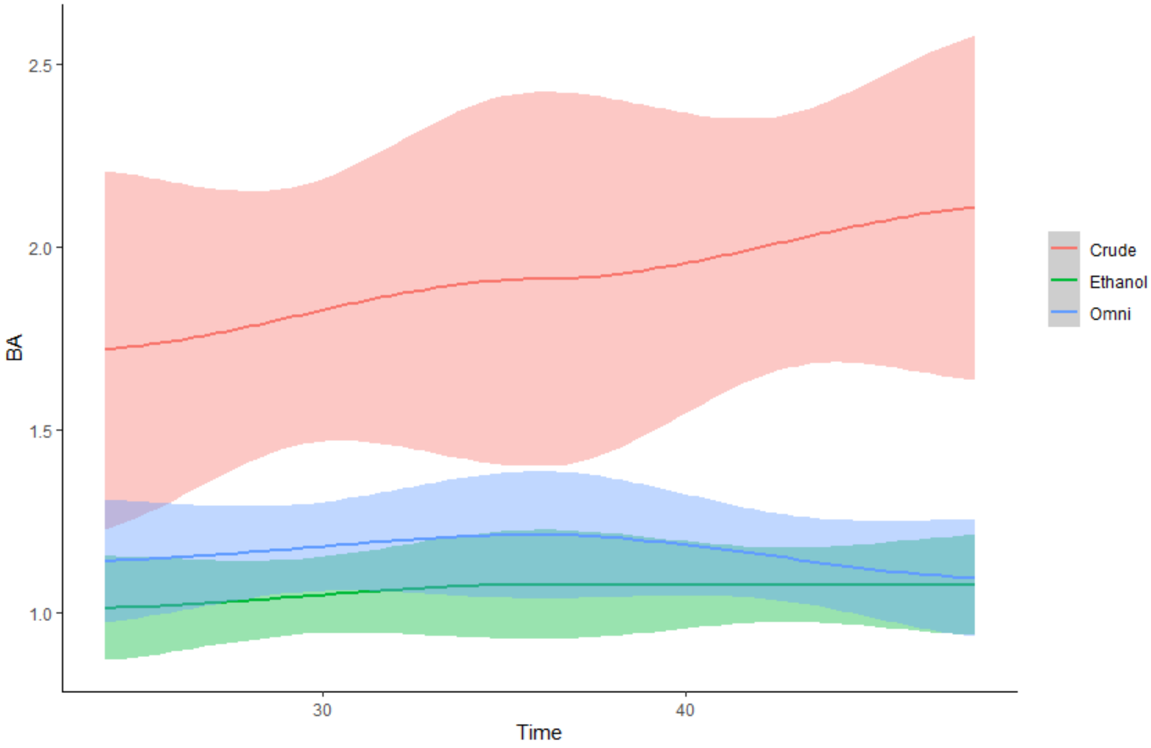

**Supplementary FigureS4.** The loess curve plot of butyric over time (24h, 36h, 48h) between stool samples collected as crude, 95% EtOH,OMNImet®•GUT .

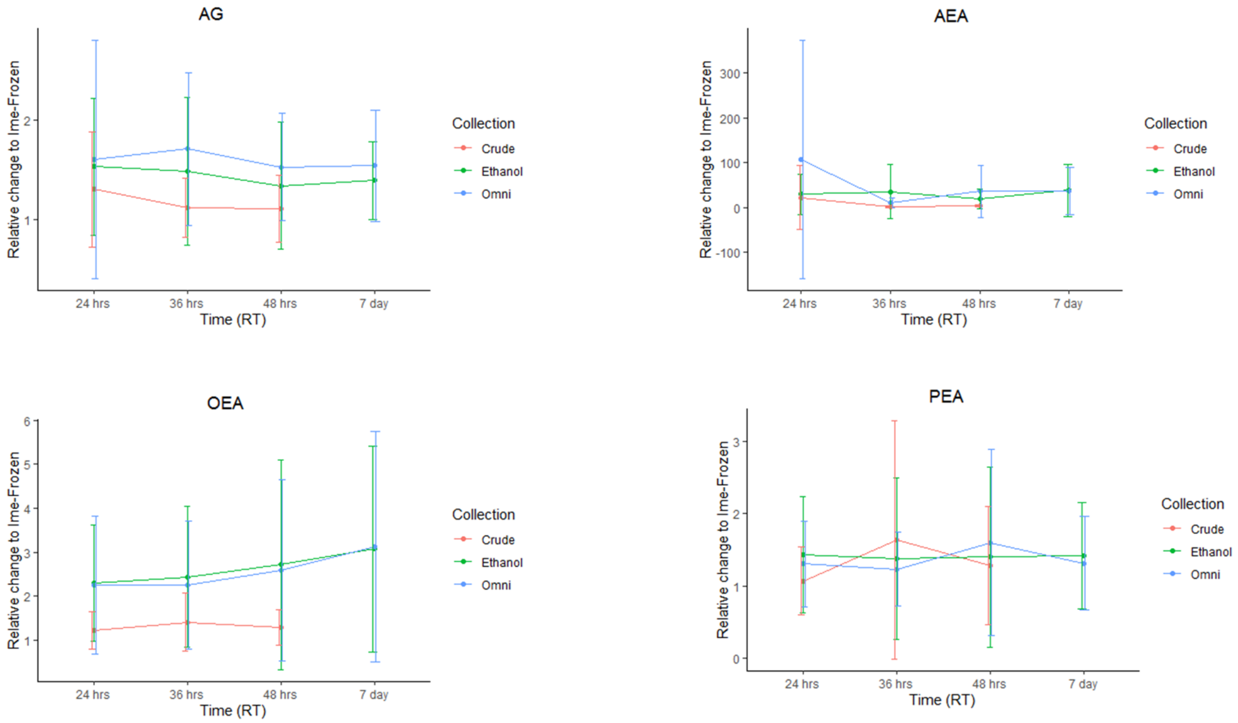

**Supplementary FigureS5** A plot showing the changes in the levels of endocannabinoids (ECCs) over time (24h, 36h, 48h, 7 day) in feces samples collected as crude, in 95% EtOH, and with OMNImet®•GUT solvent.

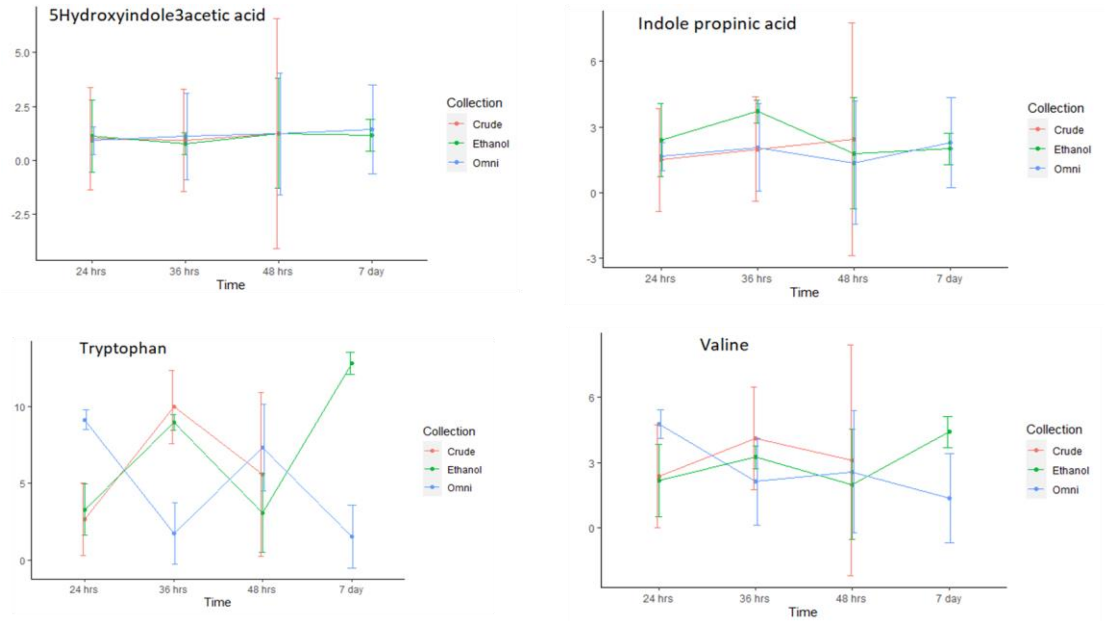

**Supplementary FigureS6** A plot showing the changes in the levels of representative class of polar metabolites changing over time (24h, 36h, 48h, 7 day) in feces samples collected as crude, in 95% EtOH, and with OMNImet®•GUT solvent.

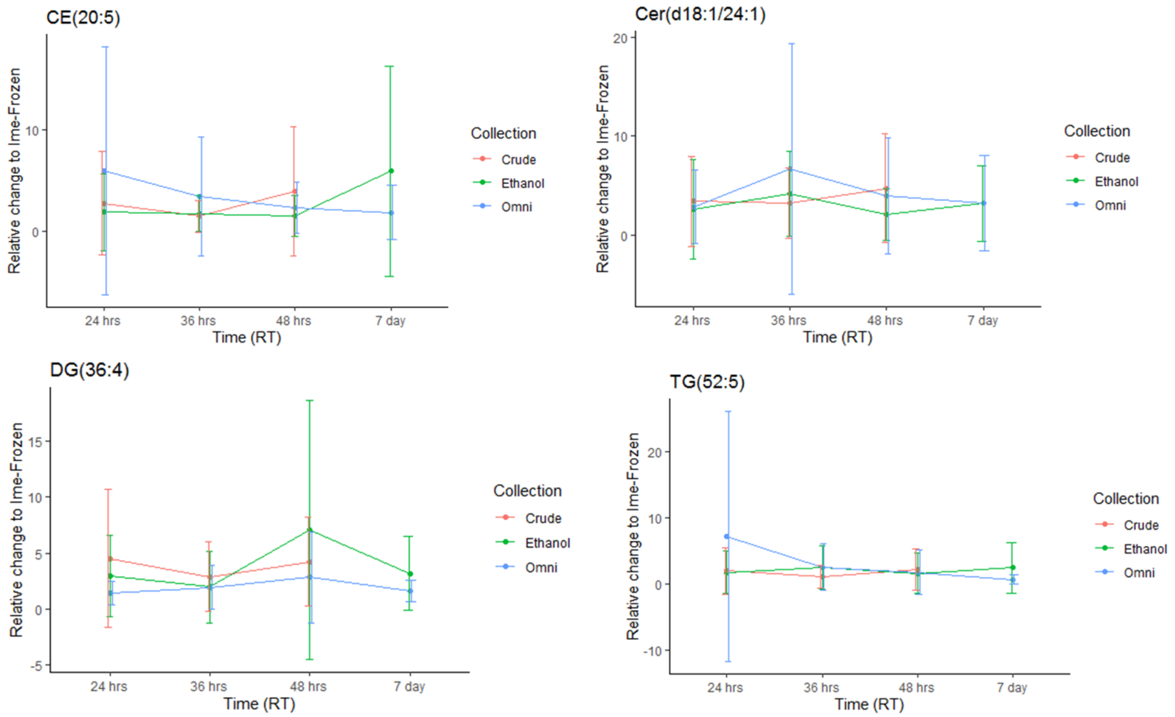

**Supplementary FigureS7** A plot showing the changes in the levels of representative class of lipids over time (24h, 36h, 48h, 7 day) in feces samples collected as crude, in 95% EtOH, and with OMNImet®•GUT solvent.

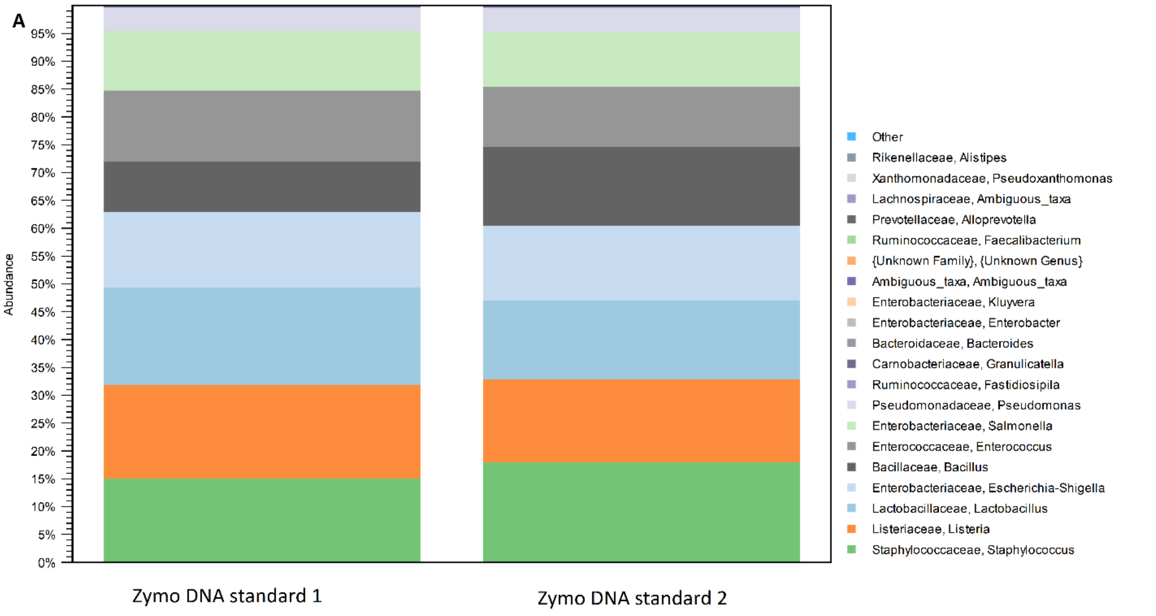

**B**

| Microbial Community DNA Standard | Theoretical abundance % | Identified level and abundance% |
|---|---|---|
| Pseudomonas aeruginosa | 4,2 | Pseudomonas 4,2% |
| Escherichia coli | 10,1 | Escherichia/shigella 14% |
| Salmonella enterica | 10,4 | Salmonella 11% |
| Lactobacillus fermentum | 10,4 | Lactobacillus 17% |
| Enterococcus faecalis | 9,9 | Enterococcus 13% |
| Staphylococcus aureus | 15,5 | Staphylococcus 15% |
| Listeria monocytogenes | 14,1 | Listeria 17% |
| Bacillus subtilis | 17,4 | Bacillus 9,1% |

Supplementary FigureS8. Positive controls A) Relative abundances of replicates of positive cotrols (DNA standard, Zymo Research) B) Table of theoretical abundance % of DNA standard and identified level and abundance %.

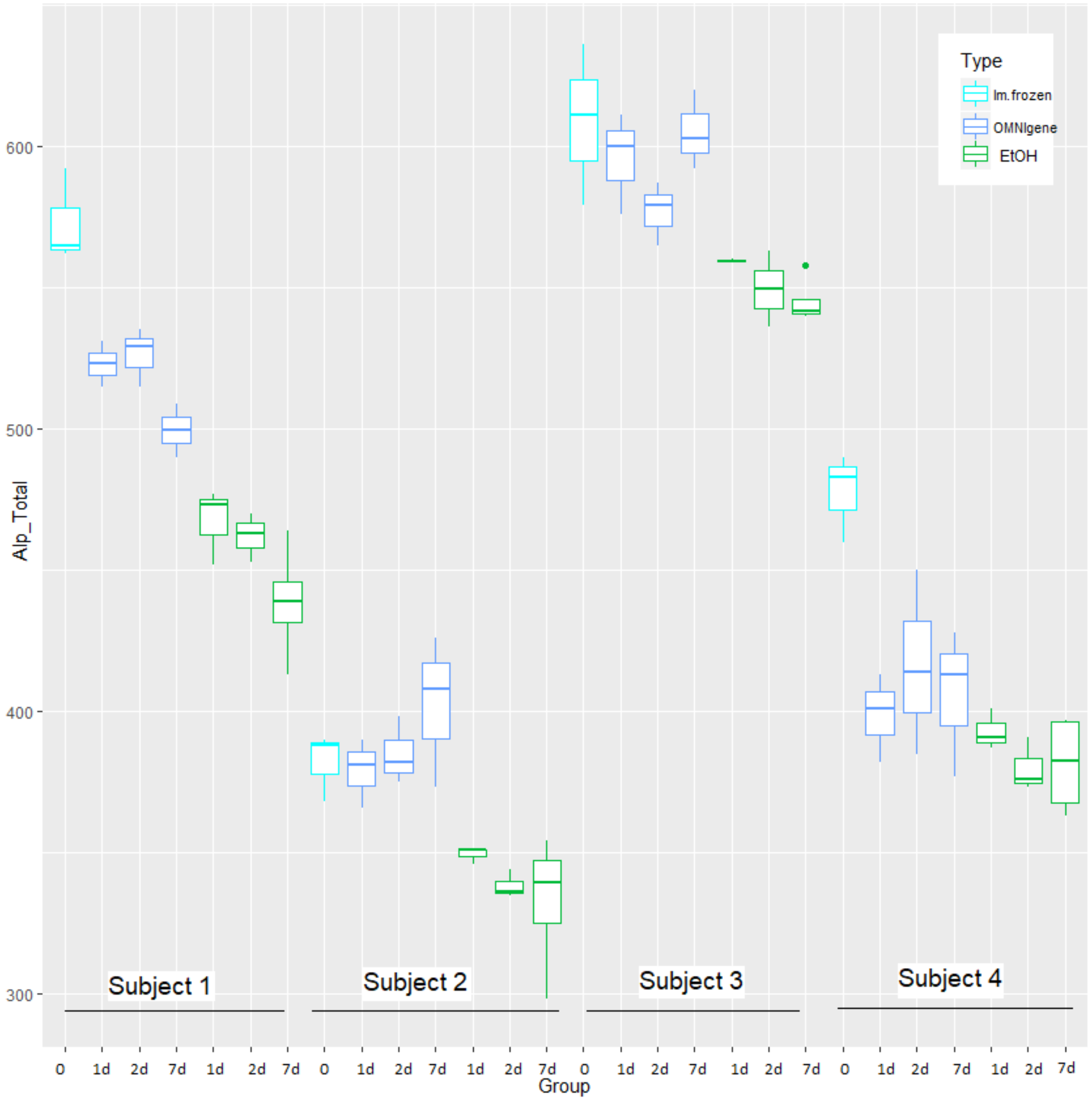

**Supplementary FigureS9** The differences in alpha diversity (Shannon index) among the three study groups.

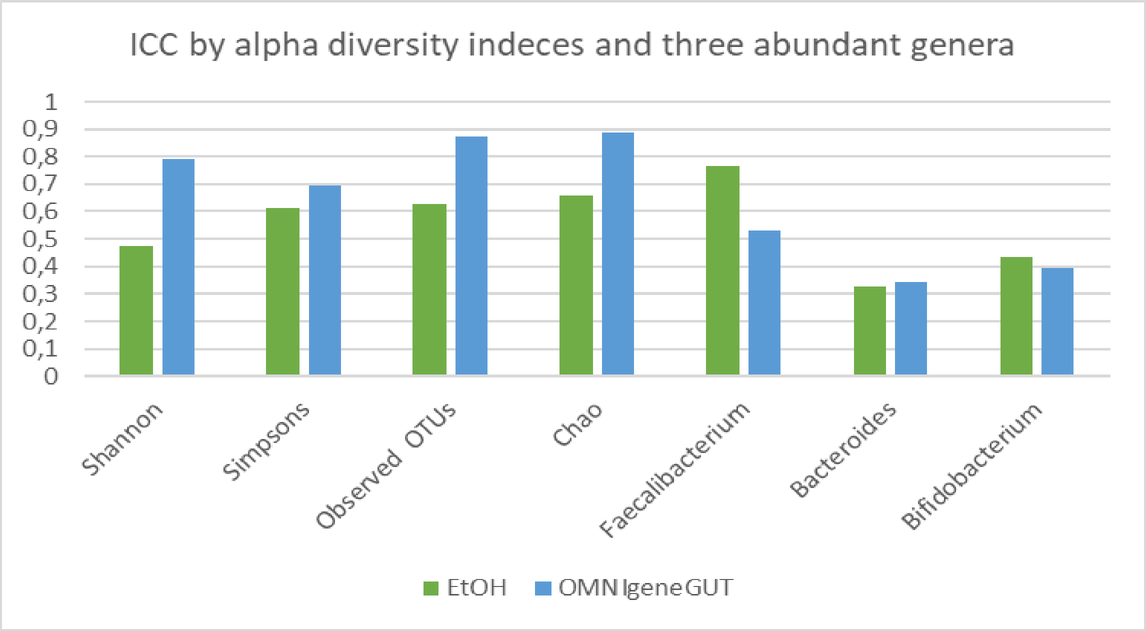

**Supplementary FigureS10** Compared similarity between storage types using intra-class correlation coefficients with immediately frozen samples used as reference.

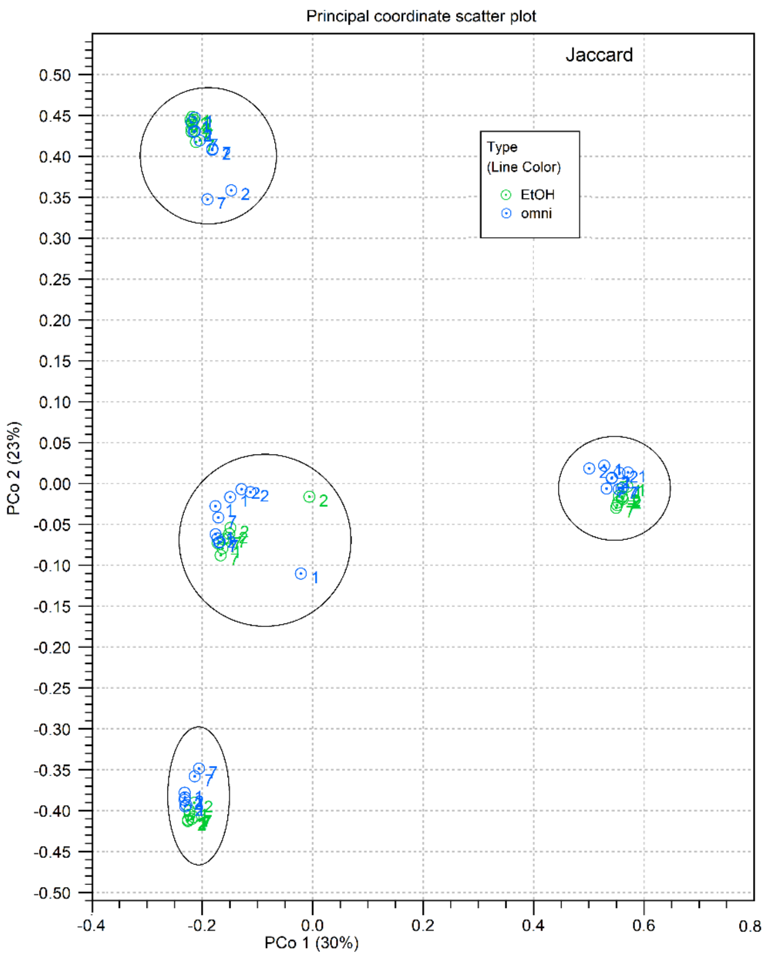

Supplementary FigureS11. Beta diversity by Principal coordinates analysis with Jaccard dissimilarity metrics. Colors indicate storage types and numbers of dots show storage days. Oval shapes form clusters of subjects.

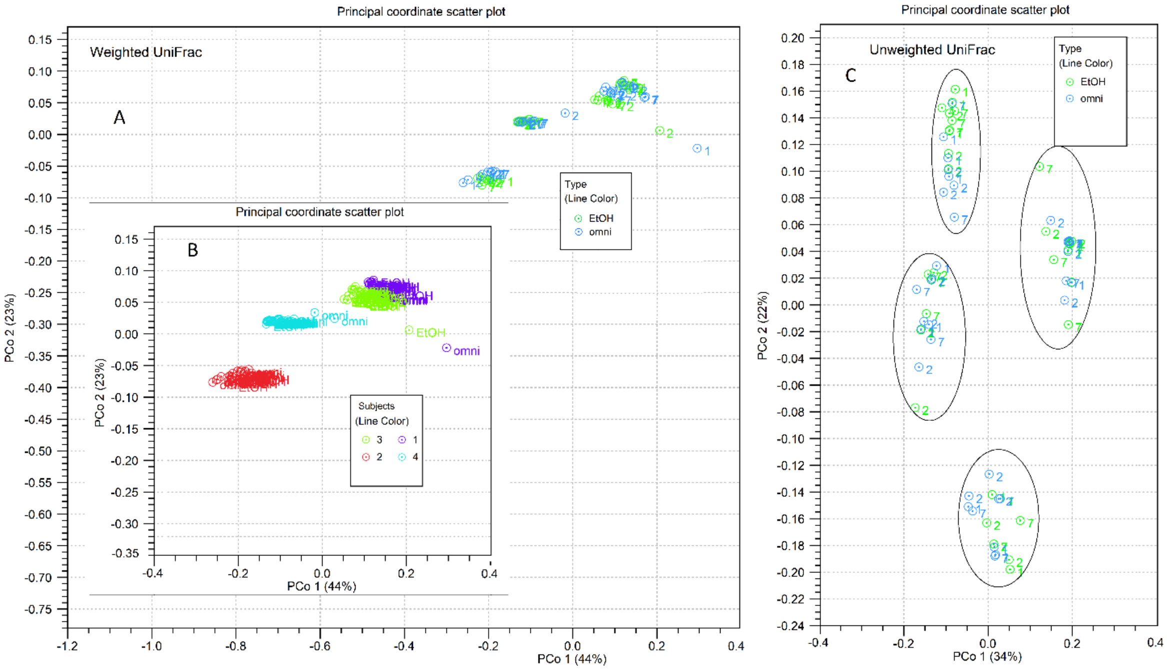

Supplementary FigureS12. Beta diversity by Principal coordinates analysis with Weighted and Unweighted UniFrac dissimilarity metrics. A) Weighted UniFrac, colors indicate storage types and numbers of dots show storage days. B) Weighted UniFrac, colors indicate different subjects C) Unweighted UniFrac, colors indicate storage types and numbers of dots show storage days. Oval shapes form clusters of subjects.
